# *Firmicutes* and *Bacteroidetes* explain mass gain variation in an obligate hibernator

**DOI:** 10.1101/2021.09.24.461421

**Authors:** Gina C. Johnson, Samuel Degregori, Paul H. Barber, Daniel T. Blumstein

## Abstract

1. Body condition is an important life history challenge that directly impacts individual fitness and is particularly important for hibernating animals, whose maintenance of adequate body fat and mass is essential for survival.
2. It is well documented that symbiotic microorganisms play a vital role in animal physiology and behaviour. Recent work demonstrates that gut microbes are associated with fat accumulation and obesity; *Firmicutes* is consistently associated with obesity while *Bacteroidetes* is associated with leanness both in humans and other animals.
3. The focus of most microbiome studies has been on human health or involved lab reared animals used as a model system. However, these microbes likely are important for individual fitness in wild populations and provide potential mechanistic insights into the adaptability and survival of wildlife.
4. Here we test whether symbiotic microorganisms within the phyla of *Firmicutes* and *Bacteroidetes* are associated with summer mass gain in an exceptionally well-studied wild population of yellow-bellied marmots (*Marmota flaviventer*) by quantifying microbial abundance over five years of fecal samples (2015 – 2019) collected during their summer active season.
5. Results show that marmots with higher mass gain rates have a greater abundance of *Firmicutes*. In contrast, higher abundance of *Bacteroidetes* was associated with lower mass gain rates, but only for marmots living in harsher environments. Similar patterns were found at the family level where *Ruminococcaceae*, a member of *Firmicutes*, was associated with higher mass gain rates, and *Muribaculaceae*, a member of *Bacteroidetes*, was associated with lower mass gain rates, and similarly in harsher environments.
6. Although correlative, these results highlight the importance of symbiotic gut microbiota to mass gain in the wild, a trait associated with survival and fitness in many taxonomic groups.

## Introduction

The maintenance of sufficient body condition is a major life history challenge shared by many animals with important implications for individual fitness (Gaillard et al., 2000; Green, 2001; Schulte-Hostedde et al., 2001). Animals in good condition can endure longer fasting periods (Atkinson & Ramsay, 1995), are more likely to survive long migrations (Merilä & Svensson, 1997), maintain a more responsive immune system (Navarro et al., 2003), have increased fecundity (Tammaru et al., 1996) and enjoy higher mating success (Cotton et al., 2006).

One important measure of body condition affecting individual survival in many taxa is relative body mass (Jakob et al., 1996; Schulte-Hostedde et al., 2001). For instance, larger body mass increases the probability of survival in bighorn sheep (*Ovis canadensis*) (Festa-Bianchet et al., 1997). Similarly, heavier canvasbacks (*Aythya valisineria*) have greater overwinter and annual survival probabilities (Haramis et al., 1986). In contrast, great tits (*Parus major*) reduced body mass in the presence of sparrowhawks (*Accipiter nisus*), reducing predation risk (Gosler et al., 1995).

While a variety of factors such as food availability, predation risk, and temperature influence individual body mass (Lima, 1986), a growing body of literature suggests that symbiotic microorganisms, collectively referred to as “microbiomes”, also play a key role in shaping host physiology (Neish, 2009; Kinross et al. 2011; Hird, 2017). The complex network of microbes that reside in the vertebrate gastrointestinal tract influences the host’s metabolic activity and affects numerous aspects of physiology, anatomy, and behaviour (Cryan & Dinan, 2012; Nicholson et al., 2012). Research on humans and other animals suggests a strong link between the intestinal microbiome and mass gain (Ley et al., 2006; Tsai & Coyle, 2009; Million et al., 2012), with shifts in the dominant phyla of gut bacteria leading to obesity (Ley et al., 2005, 2006; Turnbaugh et al. 2006, 2008; Ley, 2010).

Most gut bacteria belong to four major phyla: *Firmicutes, Bacteroidetes, Proteobacteria*, and *Actinobacteria* (Tilg & Kaser, 2011), sometimes referred to as the “core microbiome” (Turnbaugh et al. 2009; Hird et al., 2015). Studies in mammals suggest that shifts in relative abundance with more *Firmicutes* and fewer *Bacteroidetes* is associated with fat accumulation and potential for obesity; in contrast, weight loss and leanness are associated with higher relative abundance of *Bacteroidetes* (Ley et al., 2005; Turnbaugh & Gordon, 2009). The balance of *Firmicutes* and *Bacteroidetes* in the gut is often described as a ratio, *Firmicutes/Bacteroidetes* (F/B), with obese individuals having a higher F/B ratio (Eckburg et al., 2005) and lean individuals a lower F/B ratio (Mathur & Barlow, 2015; Koliada et al., 2017). That relative abundance of *Firmicutes* and *Bacteroidetes* influences obesity/leanness suggests that gut microbiomes can affect energy extraction from the diet (Ellekilde et al., 2014), with strong implications for individual fitness, especially for animals whose survival depends on developing and maintaining adequate fat stores.

Developing fat stores is critical to hibernating animals, whose long-term survival and growth depend on adequate body fat accumulation (Turbill et al., 2001). Hibernation involves dramatic seasonal changes in individual food consumption, body mass, and energy expenditure (Lyman & Chatfield, 1955; Florant et al., 2004). Moreover, hibernation alters the composition and diversity of gut microbial communities across a diversity of taxa, ranging from mammals (Dill-McFarland et al., 2014; Sonoyama et al., 2009; Malinčiová et al., 2017) to amphibians (Kohl & Yahn, 2016; Weng et al., 2016), and reptiles (Tang et al., 2019), suggesting that gut microbiota may have functional importance in hibernating animals (Carey & Assadi-Porter, 2017). For example, in juvenile Arctic ground squirrels, increases in *Bacteroidetes* and reductions of *Firmicutes* reduces individual adiposity (Stevenson, Duddleston, & Buck, 2014; Stevenson, Buck, & Duddleston, 2014) showing that microbial composition affects fat deposition. Similarly, there is variation in gut microbiomes during the hibernation and active phases of brown bears, and inoculating germ free mice with bear summer microbiota increases mice fat accumulation (Sommer et al., 2016).

Yellow-bellied marmots (*Marmota flaviventer*) are obligate hibernators that must accumulate sufficient fat stores to survive a 6-8 month hibernation (Hall & Kelson, 1959; Armitage, 1998; Armitage et al., 2003). Marmots lose up to half of their body mass during hibernation (Armitage et al., 1976), thus mass gain during the active season is essential for survival. Moreover, adequate fat stores are essential for reproductive female marmots to give birth, directly influencing individual reproductive fitness (Andersen et al., 1976). Environmental conditions largely explain variation in mass gain in marmots (Maldonado-Chaparro et al., 2015), but age, sex, diet, food availability, and body size can also play a role (Armitage et al., 1976, 2003). However, given seasonal shifts in gut microbiomes of other hibernating animals (Stevenson et al., 2014a; Stevenson et al., 2014b; Sommer et al., 2016) and that microbiome composition can influence mass gain (Ley et al., 2005; Turnbaugh & Gordon, 2009; Ellekilde et al., 2014), it is possible that gut microbiome composition could influence the survival and reproductive fitness of individual marmots.

We examined the association of symbiotic gut bacteria and mass gain rate in an exceptionally well-studied wild population of yellow-bellied marmots. This population has been continuously monitored since 1962, providing long-term data on mass gain during the active season, overwinter and summer survival, and reproductive success. Data shows that age and sex play a critical role in mass gain (Armitage, 2014), with factors like chronic stress and spatiotemporal variation affecting individual survival (Ozgul et al., 2006; Wey et al., 2014). Here we use 16S microbial metabarcoding to examine microbial composition in free-living marmots to estimate the relative effects of microbiome composition and environmental factors in explaining variation in mass gain rates. Specifically, we test the hypothesis that *Firmicutes* abundance is associated with greater mass gain while *Bacteroidetes* abundance is associated with lower mass gain rates.

## Materials & Methods

### Study species and site

We studied yellow-bellied marmots in and around the Rocky Mountain Biological Laboratory (RMBL), located in the Upper East River Valley in Gothic, Colorado, U.S.A. (38°77′N, 106°59′W). We trapped marmots by placing Tomahawk-live traps near burrow entrances. After capture, the marmots were transferred to cloth handling bags to measure their body mass (to the nearest 10 g), and to determine their sex and reproductive status (Blumstein et al., 2006). Each marmot was given a set of unique ear-tag numbers and their dorsal pelage marked with Nyanzol fur dye for identification from afar. Faecal samples are easily collected throughout the season when animals are live trapped. When faeces are found in traps, they are routinely collected in a plastic bag, immediately put on ice, and subsequently frozen at -20 °C within 2 h of collection. Samples are then transported from the field on dry ice and stored at -80 °C in the lab for long-term preservation.

From a large archive of faecal samples, we selected paired samples from individual females to capture variation within the active season, although for some individuals only a single sample was available. We focused on females, because both overwinter survival and reproduction the next year depends on body condition (Andersen et al., 1976). We collected samples from 10 different colonies: 5 from higher elevation colonies (mean elevation 3,043 m) and 5 lower elevation colonies (mean elevation 2,883 m), separated by a maximum horizontal distance of 4.9 km. Although there is only an average of 160 m in altitude between these sites, the phenology of these locations differs substantially, resulting in emergence from hibernation and mating approximately 2 weeks earlier in down-valley colonies (Blumstein, 2009).

We selected samples collected closest to 1 June and 15 August, dates that fall within the period of linear mass gain during their active season (Heissenberger et al., 2020), although some samples were collected as early as May and as late as September. To maximise statistical power and test for consistency across time, we selected samples across a five year span (2015 – 2019), yielding 207 total samples representing 71 individuals. We sampled multiple age groups, 25 juvenile, 67 yearlings, and 109 adults (Table 1) because each age class faces unique ecological challenges (Armitage et al., 1976; Heissenberger et al., 2020). Juveniles emerge from their natal burrow in late June early July and must rapidly gain mass and body size to survive their first hibernation, despite not reaching full body size until their second year. Yearlings tend to show the largest change in mass as they gain fat to survive hibernation but also undergo somatic growth to reach adult body size during their second summer active season. Adults (defined as reproductively mature females in their third year of life or older) have typically reached full body size so only need to accumulate sufficient fat stores during their summer active season (Armitage et al., 1976).

**Table 1.** Characteristics of yellow-bellied marmots among selected samples (*N* = 201)

| Characteristics | Number of samples (unique individuals) |  |  |  |  |
| --- | --- | --- | --- | --- | --- |
| Year | 2015 | 2016 | 2017 | 2018 | 2019 |
| Age Class |  |  |  |  |  |
| *Adult* | 11 (6) | 13 (8) | 26 (13) | 22 (11) | 37 (20) |
| *Yearling* | 4 (3) | 20 (11) | 8 (4) | 9 (5) | 26 (14) |
| *Juvenile* | 2 (1) | 3 (2) | 7 (4) | 0 | 13 (7) |
| Valley Position |  |  |  |  |  |
| *Up-Valley* | 16 (9) | 29 (16) | 30 (15) | 20 (10) | 27 (14) |
| *Down-Valley* | 1 (1) | 7 (5) | 11 (6) | 11 (6) | 49 (27) |

### Microbiome sample processing and sequencing

We isolated bacterial DNA from faecal samples with Qiagen Powersoil Extraction kits following the manufacturer’s protocol (Germantown, Maryland, USA). We generated 16S libraries using the 515F (5’-GTGCCAGCMGCCGCGGTAA) and 806R (5’-GGACTACHVHHHTWTCTAAT) primers targeting the V4 region of the 16S rRNA gene (Caporaso et al., 2011). Samples underwent PCR, in triplicate 25 ul reactions, using a Qiagen Multiplex PCR kit with the following thermocycler conditions: 1 cycle of 94 °C for 3 min; 35 cycles of 94 °C for 45 s, 50 °C for 60 s and 72 °C for 90 s; and 1 cycle of 72 °C for 10 min (Thompson et al., 2017). We pooled triplicate reactions after confirming amplification success through gel electrophoresis, and then dual-indexed samples using the Nextera UD Index Kit (Ilumina, San Diego, USA) and then purified with OMEGA Bio-Tek MagBind magnetic beads (Norcross, Georgia, USA). Laragen (Culver City, California, USA) performed quantification and pooling to create libraries with equimolar sample concentrations. Multiplexed libraries were paired-end sequenced (300 bp per sequence) on an Ilumina Miseq v3 at Laragen. We carried negative controls from the DNA extraction process and subsequent PCR’s throughout sample processing and added these to the final pooled library for sequencing.

### Data quality control and analysis

The resulting sequence libraries were run through the QIIME2 (v. 2019.9) microbiome data science platform (Bolyen et al., 2018) for quality control, amplicon sequence variant (ASV) taxonomy assignment, and community diversity analyses. Data were demultiplexed and denoised using dada2 (Callahan et al., 2016) and merged into a feature table for analysis. We then rarefied samples to a minimum sequencing depth of 1000 reads, all samples with fewer than 1000 reads were excluded from analysis. ASVs were assigned taxonomy using a naive Bayes taxonomy classifier trained on the SILVA database (Quast et al., 2013; Yilmaz et al., 2014; Glöckner et al., 2017) with reference sequences clustering at 99% similarity. ASVs with fewer than 5 reads were pruned as well as ASVs occurring in less than 3% of the samples (Karstens et al., 2019). Any ASVs associated with assignments to eukaryotes, chloroplasts, and cyanobacterial reads were also pruned. ASVs were compiled into a table and analyzed in R version 3.5.1. (R Core Team, 2017) using the package *phyloseq* (McMurdie & Holmes, 2013).

### Estimating mass gain rate

Because marmot mass gain rates vary with age (Armitage et al., 1976), repeated measures of body mass were taken for all individuals captured from 2015 – 2019. Using methods from Heissenberger et al. (2020), we used body mass at emergence from the natal burrow for juveniles, predicted 1 June body mass for yearlings and adults, and 15 August body mass for all ages, dates that reflect the bulk of the growing season for each respective age class. Predicted values were calculated by fitting a linear mixed effects model on body mass measurements, where individual identity, year, and site were included as random effects and colony, age and sex were fixed effects. This permitted us to generate Best Linear Unbiased Predictions (BLUPS) for predicting 1 June and 15 August body mass (Heissenberger et al., 2020; Maldonado-Chaparro et al., 2015; Ozgul et al., 2010). We calculated juvenile growth rate as the difference from the 15 August body mass and the mass at first natal emergence divided by the number of days between them. For yearlings and adults, it was the difference between 15 August and 1 June masses divided by the number of days between them (76 d).

### Testing bacterial composition influence on mass gain

We fitted linear mixed effects models (Bates et al., 2015) to explain variation in mass gain rates. To account for limitations in sequencing and to control for spurious correlations, the phyla OTU tables were transformed using the centered log ratio, or CLR (Aitchison, 1986; Gloor et al., 2017). Models included the fixed effects of CLR transformed ASV counts assigned to *Bacteroidetes* or *Firmicutes*, and subsequent families within those phyla (these continuous variables were zero and centered), valley position, and age class, and the interactions between bacterial phyla or family and valley position and bacterial phyla or family and age class. We included year and marmot ID as random effects. We included valley position, because snow melts later at the higher elevation sites and this potential for an effect of elevation on mass gain, combined with later marmot emergence means that they live in a relatively harsher environment with less time to gain mass (Vuren & Armitage, 1991; Blumstein, 2009; Armitage, 2014). Non-significant interactions were removed and the models refitted for interpretation (Engqvist, 2005). We estimated the marginal and conditional R^2^ values using the package *MuMIn* (Bartoń, 2015). Lastly, we estimated the relative amount of variation explained by the bacterial taxa by removing either the bacterial taxa or the significant interaction between the bacterial taxa and another fixed effect, and refitting the final model without it.

## Results

We amplified 207 samples, yielding a total of 10,930,721 raw sequencing reads. After cleaning and filtering, 2,449,899 reads remained and were merged into a feature table for analysis. Sample sequencing depth ranged from 27 reads to 71,502 reads. As such, we rarefied all samples to a minimum depth of 1000 reads, excluding a total of 6 samples with fewer than 1000 reads from subsequent analysis, leaving a total of 201 samples representing 71 individuals across five years (2015 - 2019).

### Bacterial taxonomic composition of the marmot gut microbiome

At the phylum level, *Firmicutes* dominated marmot gut microbiomes, averaging 61% abundance across all samples. *Bacteroidetes* was the second most dominant group, averaging 29% followed by *Tenericutes* with an average of 5.6% across all samples. *Actinobacteria, Proteobacteria, and Verrucomicrobia* occupied the rest of most samples; the presence of other groups was less than 1% (Figure 1A). Examining microbial abundance at the family level showed that *Ruminococcaceae* was the most dominant with an average abundance of 35%, followed by *Lachnospiraceae* with 15%, *Muribaculaceae* with 12%, and *Rikenellaceae* with 8.6% mean abundance across all samples. *Rikenellaceae, Bacteroidaceae, Christensenellaceae, Clostridiales vadinBB60 group, Anaeroplasmataceae*, and *Erysipelotrichaceae* occupied the rest of most samples (Figure 1B). *Ruminococcaceae* and *Lachnospiraceae* families are members of the phylum *Firmicutes*, while *Muribaculaceae* and *Rikenellaceae* families are part of the *Bacteroidetes* phylum.

**Figure 1.**
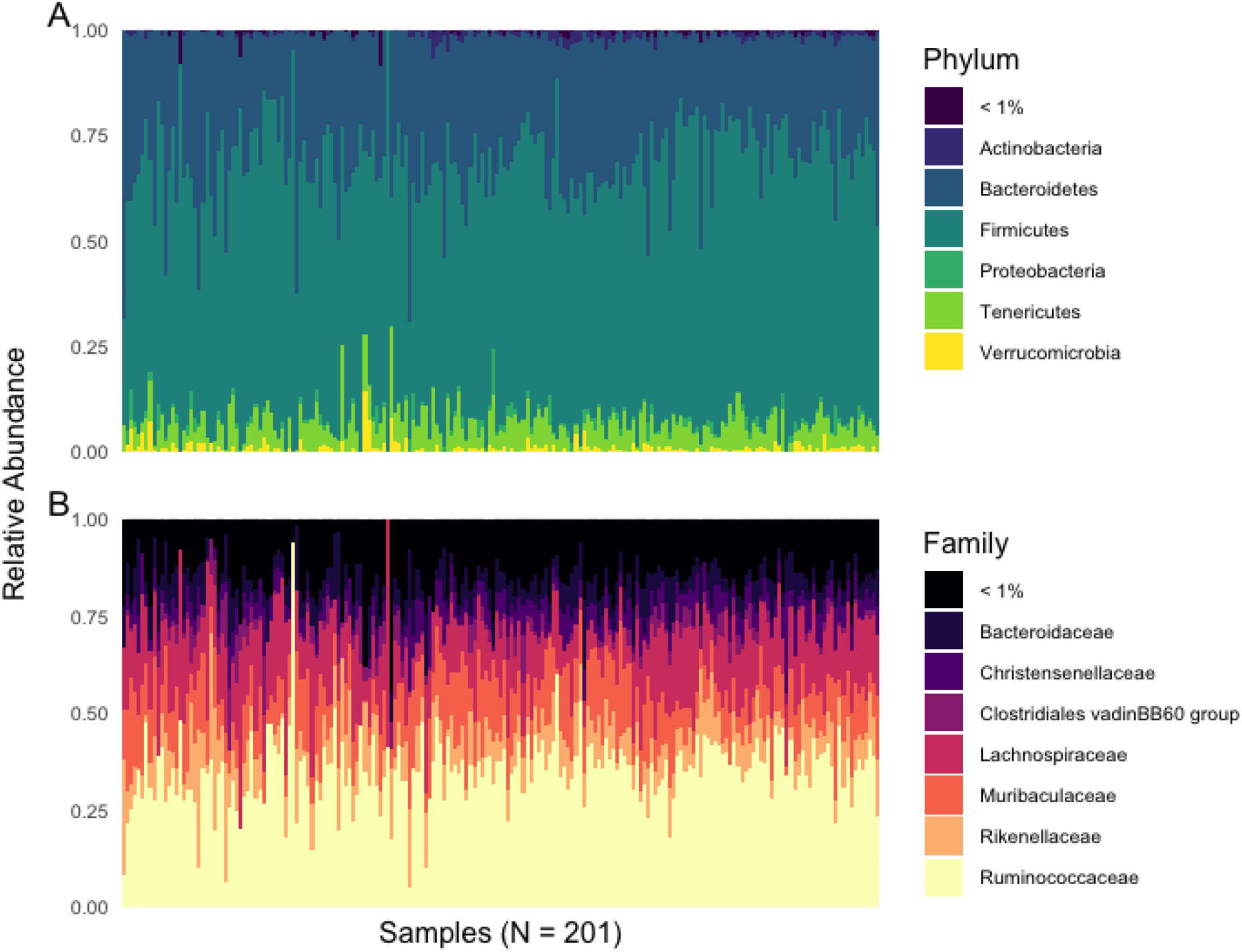
A) The relative abundance of dominant gut phyla across all samples (*N* = 201) showing *Firmicutes* and *Bacteroidetes* occupying the majority of reads. B) The relative abundance of dominant gut families across all samples (*N* = 201).

### Gut microbes explain variation in marmot mass gain rates

Mass gain rates across all 71 individuals varied by age class. Adults gained mass at an average of 15.29 g/day, yearlings 21.08 g/day, and juveniles 24.53 g/day. The distribution of mass gain rates conformed to normal expectations (*W* = 0.99185; *P* = 0.3223) and therefore was not transformed. After controlling for variation explained by age class and valley position as fixed effects, and year and individual identity as random effects, we found that abundance of *Firmicutes* was positively associated with variation in mass gain rates (Figure 2; Table 2; estimate = 0.645 ± 0.237 sem; *P* = 0.007; estimated partial *R*^*2*^ = 0.011). *Bacteroidetes* was negatively associated with variation in mass gain rates although with the opposite relationship, and only in up valley colonies (Figure 3; Table 3; estimate = -0.974 ± 0.432 sem; *P* = 0.026; estimated partial *R*^*2*^ = 0.039).

**Table 2.** Fixed and random effects from the best fit model showing *Firmicutes* and age class explain variation in mass gain rates. The adult age class is used as the reference category. Significant effects are shown in bold.

| Variable | Estimate (SE) | *t* | *P* |
| --- | --- | --- | --- |
| **(Intercept)** | **14.693** | **10.107** | **1.9e<sup>-07</sup>** |
| ***Firmicutes*** | **0.645** | **2.712** | **0.007** |
| Valley Position | 1.601 | 1.437 | 0.155 |
| **Age Class (J)** | **-3.505** | **-3.101** | **0.002** |
| **Age Class (Y)** | **4.499** | **6.749** | **1.67e<sup>-10</sup>** |
| Random Effects |  |  |  |
| Groups Name | Variance | Std. Dev. |  |
| Individual UID | 18.229 | 4.270 |  |
| Year | 5.464 | 2.337 |  |
| Observations | 201 |  |  |
| Marginal $R^2$ / Conditional $R^2$ | 0.175 / 0.831 | | |

**Table 3.** Fixed and random effects from the best fit model showing *Bacteroidetes* in up valley colonies (UV) and age class explaining variation in mass gain rates. The adult age class is used as the reference category. Significant effects are shown in bold.

| Variable | Estimate (SE) | *t* | *P* |
| --- | --- | --- | --- |
| **(Intercept)** | **15.008** | **10.675** | **3.98e<sup>-08</sup>** |
| *Bacteroidetes* | 0.092 | 0.293 | 0.770 |
| Valley Position (UV) | 1.167 | 1.052 | 0.296 |
| **Age Class (J)** | **-3.158** | **-2.763** | **0.006** |
| **Age Class (Y)** | **4.380** | **6.581** | **4.24e<sup>-10</sup>** |
| ***Bacteroidetes*\*UV** | **-0.975** | **-2.253** | **0.026** |
| Random Effects |  |  |  |
| Groups Name | Variance | Std. Dev. |  |
| Individual UID | 18.406 | 4.290 |  |
| Year | 4.878 | 2.209 |  |
| Observations | 201 |  |  |
| Marginal $R^2$ / Conditional $R^2$ | 0.172 / 0.829 | | |

**Figure 2.**
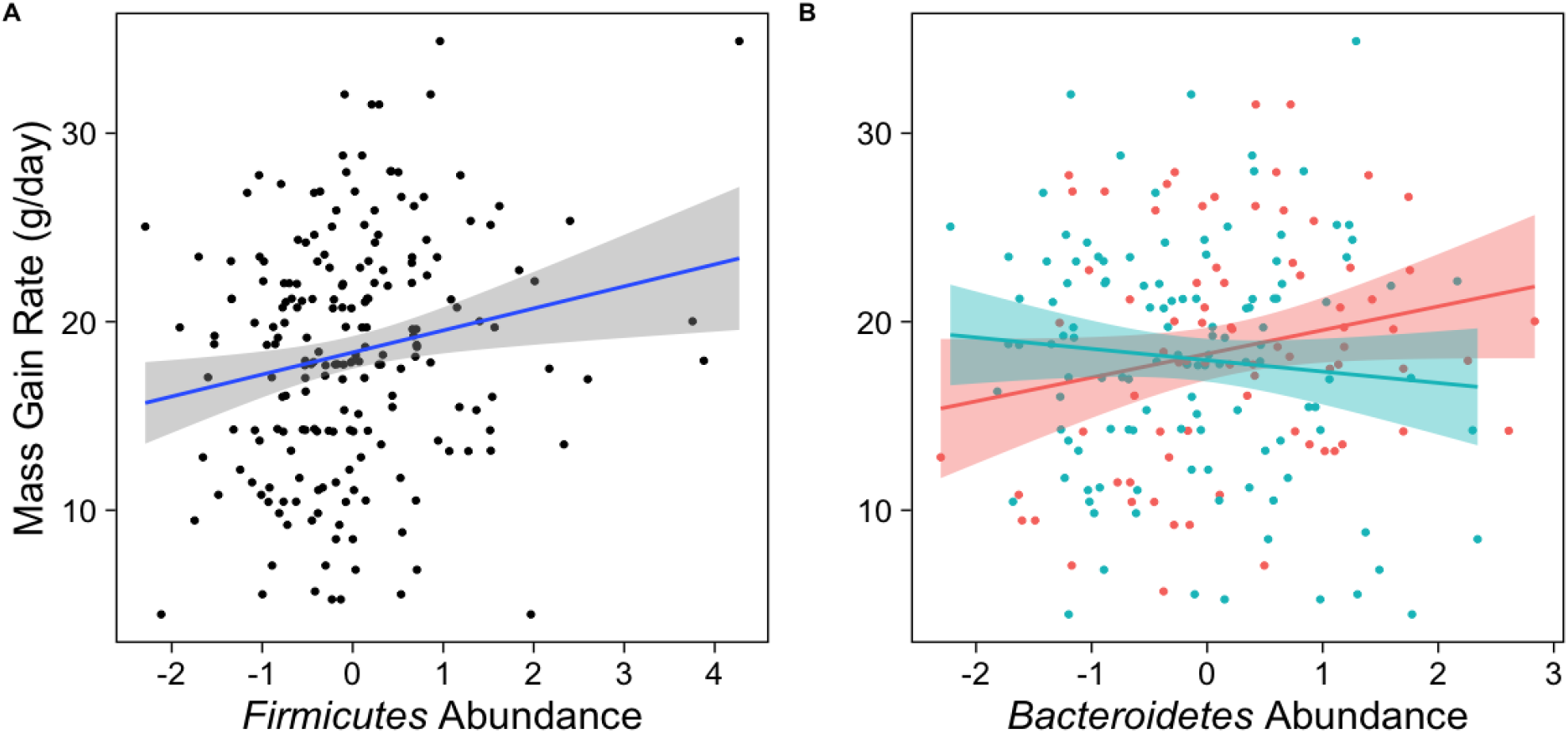
A) Relationship between *Firmicutes* abundance and mass gain rate for all samples (*N* = 201) across the marmot active season. The blue line shows the predicted relationship based on the linear mixed effects model. B) Relationship between *Bacteroidetes* abundance and mass gain rate for all samples (*N* = 201) across the marmot active season. The blue line shows the predicted relationship based on the linear mixed effects model between *Bacteroidetes* abundance mass gain rate in higher elevation colonies, while the red line shows the predicted relationship between *Bacteroidetes* abundance and mass gain rate in lower elevation colonies.

**Figure 3.**
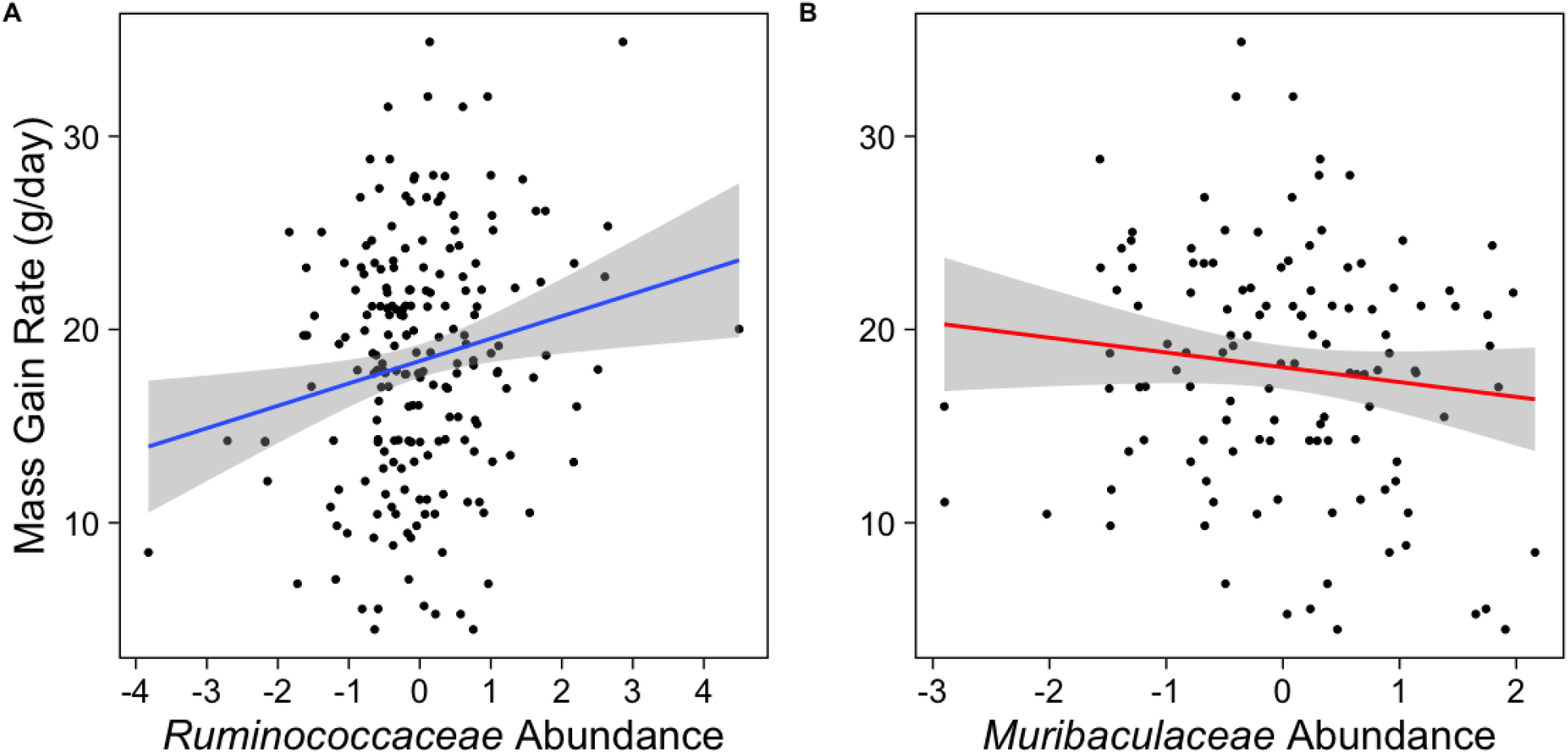
A) Relationship between *Ruminococcaceae* abundance and mass gain rate for all samples (*N* = 201) across the marmot active season. The blue line shows the predicted relationship based on the linear mixed effects model. B) Relationship between *Muribaculaceae* abundance and mass gain rate for only up valley samples (*N* = 122) across the marmot active season. The blue line shows the predicted relationship based on the linear mixed effects model between *Muribaculaceae* abundance mass gain rate in higher elevation colonies.

Age class was included as a fixed effect in each mixed model, and was significantly associated with mass gain rates for both juveniles (Table 2 (*Firmicutes*); estimate = -3.505 ± 1.174 sem; *P* = 0.002)(Table 3 (*Bacteroidetes*); estimate = -3.158 ± 1.188 sem; *P* = 0.006) and for yearlings (Table 2; estimate = 4.499 ± 0.681 sem; *P* = 1.67 × 10^−10^)(Table 3; estimate = 4.380 ± 0.679 sem; *P* = 4.24 × 10^−10^). Age class explained much of the variation in mass gain rates in both the *Firmicutes* model (estimated partial *R*^*2*^ = 0.159) and the *Bacteroidetes* model (estimated partial *R*^*2*^ = 0.153). Adults were used as the reference age class. Valley position did not explain variation in mass gain rates in either of the two models (Table 2; estimate = 1.601 ± 1.209 sem; *P* = 0.155)(Table 3; estimate = 1.167 ± 1.204 sem; *P* = 0.296).

Further analysis at the family level revealed that abundance of *Ruminococcaceae* within the phylum *Firmicutes*, and *Muribaculaceae* within the phylum *Bacteroidetes* significantly explained variation in mass gain rates. After controlling for variation explained by age class, and valley position as fixed effects and year and individual identity as random effects, we found that abundance of *Ruminococcaceae* was positively associated with variation in mass gain rates (Figure 3A ; Table 4 ; estimate = 0.528 ± 1.<u>496 sem</u>; *P* = 0.016; estimated partial *R*^*2*^ = 0.017). *Muribaculaceae* was negatively associated with variation in mass gain rates although with the opposite relationship, and only in up valley colonies (Figure 3B ; Table 5 ; estimate = -1.416 ± 0.531 sem; *P* = 0.008; estimated partial *R*^*2*^ = 0.024). We evaluated assumptions of the final models by plotting their residuals (they were approximately normal), plotting q-q plots (they were roughly straight), and plotting residuals vs. fitted values (there were no obvious patterns).

**Table 4.** Fixed and random effects from the best fit model showing *Ruminococcaceae* and age class explain variation in mass gain rates. The adult age class is used as the reference category. Significant effects are shown in bold.

| Variable | Estimate (SE) | *t* | *P* |
| --- | --- | --- | --- |
| **(Intercept)** | **14.989** | **10.017** | **4.68e<sup>-08</sup>** |
| ***Ruminococcaceae*** | **0.528** | **2.245** | **0.016** |
| Valley Position | 1.265 | 1.040 | 0.301 |
| **Age Class (J)** | **5.835** | **4.956** | **1.68e<sup>-06</sup>** |
| **Age Class (Y)** | **4.284** | **6.293** | **2.00e<sup>-09</sup>** |
| Random Effects |  |  |  |
| Groups Name | Variance | Std. Dev. |  |
| Individual UID | 18.550 | 4.307 |  |
| Year | 5.079 | 2.466 |  |
| Observations | 201 |  |  |
| Marginal $R^2$ / Conditional $R^2$ | 0.170 / 0.830 | | |

**Table 5.**
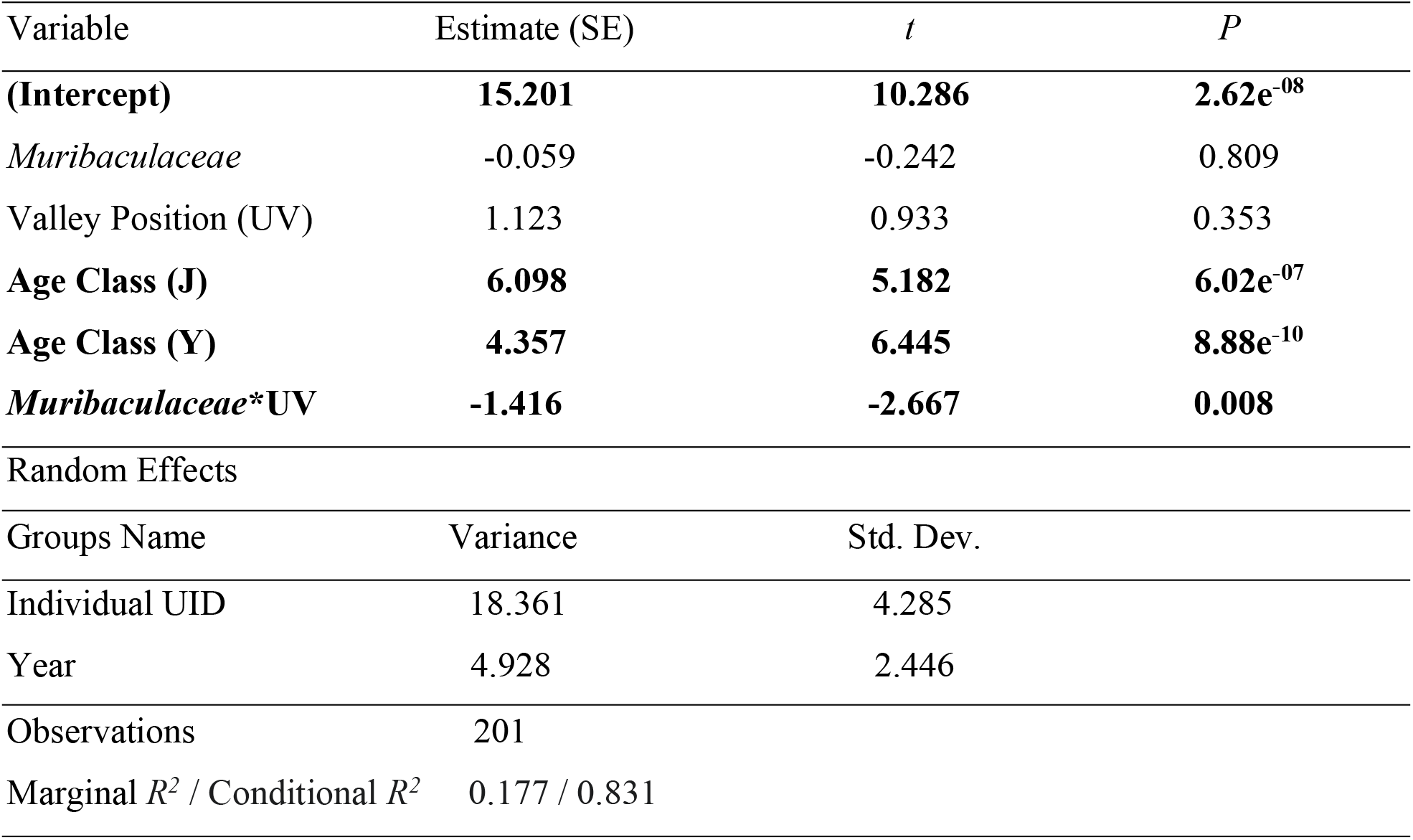
Fixed and random effects from the best fit model showing *Muribaculaceae* in up valley colonies (UV) and age class explaining variation in mass gain rates. The adult age class is used as the reference category. Significant effects are shown in bold.

Age class was also associated with mass gain rates at the family level analysis for both juveniles (Table 4 (*Ruminococcaceae*); estimate = 5.835 ± 1.177 sem; *P* = 1.68 × 10^−6^)(Table 5 (*Muribaculaceae*); estimate = 6.098 ± 1.176 sem; *P* = 6.02 × 10^−7^) and for yearlings (Table 4; estimate = 4.284 ± 0.680 sem; *P* = 2.00 × 10^−9^)(Table 5; estimate = 4.357 ± 0.676 sem; *P* = 8.88 × 10^−10^). Age class also explained much of the variation in mass gain rates in both the *Ruminococcaceae* model (estimated partial *R*^*2*^ = 0.155) and the *Muribaculaceae* model (estimated partial *R*^*2*^ = 0.157). Adults were used as the reference age class. Valley position did not explain variation in mass gain rates in either of the two models (Table 4; estimate = 1.265 ± 1.216 sem; *P* = 0.301)(Table **5**; estimate = 1.123 ± 1.204 sem; *P* = 0.353).

### Microbial composition is not influenced by age class or valley position

Individuals within the same age class or living in the same part of the valley (either at higher or lower elevation sites) did not cluster by gut microbiome composition similarities. Principle-coordinate analysis (PCoA) of Bray-Curtis distance metrics on the overall composition of the marmot fecal microbiome revealed no pattern of clustering between different age classes or individuals living up or down valley. *PERMANOVA* analysis confirmed no significant pattern across age classes (*P* = 0.162) or valley position (*P* = 0.490)(Figure S1).

## Discussion

Comparison of microbiome composition shows that relative abundance of *Firmicutes* and *Bacteroidetes* was significantly associated with mass gain in marmots, as reported for other mammals (Ley et al., 2005; Turnbaugh & Gordon, 2009). Other studies on microbiomes and mass gain remove animals from their native habitat, potentially altering the microbiome due to changes in diet, environmental factors, and interactions with humans during captivity (Uenishi et al., 2007; Dhanasiri et al., 2011; Clayton et al., 2016). Our study is unique in that we demonstrate the effects of microbiomes on mass gain in wild, free-living population.

To our knowledge, this is the first study to investigate the effect of gut microbes on mass gain in a hibernating animal from a wild population. Previous studies on hibernating species found no significant effect of gut microbe abundance on seasonal fattening (Stevenson, Duddleston, & Buck, 2014; Sommer et al., 2016), a result that may result from low sample sizes (*N* = 46 and *N* = 16) necessitated by keeping animals in captivity. By examining wild populations, our study had a much larger sample size (*N* = 201), increasing the statistical power to detect the subtle, but significant impact of microbiome composition on mass gain.

It is important to note that the effect of *Bacteroidetes* and *Firmicutes*, and the families within those phyla, *Ruminococcaceae* and *Muribaculaceae*, was comparatively small compared to other factors like age. However, this small effect size may be biologically consequential, as even a few days variation in emergence time can be the difference between survival and death (Armitage, 1976; Van Vuren & Armitage, 1991). Further investigation is needed to determine direct effects of microbes on survival. While age is a major factor (Armitage, 2014), valley position and environmental conditions (Vuren & Armitage, 1991) also explain variation in over-winter survival. Given our prior research on marmots that have shown age and sex explain much of the variation in mass gain and ability to fatten prior to hibernation, we expected the effects of microbes to be relatively small.

Like previous studies (Ley et al., 2005; Turnbaugh & Gordon, 2009), our results from marmots showed that relative abundance of *Firmicutes* and *Bacteroidetes* had a significant impact on mass gain. However, we were able to show that these patterns largely resulted from the abundance of *Ruminococcaceae* and *Muribaculaceae*. Specifically, animals with higher mass gain rates had a higher abundance of *Ruminococcaceae* (a *Firmicutes*), while animals with lower mass gain rates had a higher abundance of *Muribaculaceae* (a *Bacteroidetes*). The latter result, however, was only true in up-valley colonies where winter conditions are harsher.

While an animals’ environment can directly influence phenotype, such as white coats in coyotes and snowshoe hares during winter months, bacterial symbionts also influence phenotype (Lynch & Hsiao, 2019). Therefore, the host – microbiome – environment relationship can be complex and vary across individuals that live in different places (Koskella & Bergelson, 2020). In this study, the association of *Bacteroidetes/ Muribaculaceae* was associated with lower mass gain rates only in animals from higher elevation colonies. It could be that marmots that live in lower elevation and less harsh conditions are less likely to be influenced by variation in *Bacteroidetes* because they are able to offset the cost of having more *Bacteroidetes* by eating and gaining mass for a longer period of time. Snow melts at our lower elevation sites about two weeks earlier than our higher elevation sites leading to an extended growing season (Van Vuren & Armitage, 1991; Blumstein, 2009; Armitage, 2014). Therefore, animals living in harsher environments may be more effected by the *Bacteroidetes* than those living in less harsh conditions.

Interestingly, our results show no patterns of similarities or clustering between the different age classes or animals living at different elevations. Lack of clustering between age classes may be due to the social behavior of this species (Armitage, 1991), because regardless of age, all animals in groups within colonies are sharing the same burrow and consistently interacting with one another (Blumstein et al. 2004) and social behavior has been shown to be a direct mode for microbial transmission (Archie & Tung, 2015; Sarkar et al. 2020). Additionally, studies have shown social species to share similar microbiomes with individuals whom they have the most interactions (Moeller et al. 2016). Lack of clustering by colony elevation may be explained by similar diets in each region. The furthest distance between up and down-valley colonies is 4.9 km and vegetation types are essentially identical. It is expected that with increasing physical distance between hosts, β-diversity between hosts or groups would increase because microbial transmission is attenuated (Moeller et al. 2017). Although dispersal between these valley areas is rare, animals occupy the same valley, and are therefore not geographically separated (Armitage, 1991).

Our results are consistent with many human studies that show an association between *Firmicutes* and obesity, while leanness is associated with *Bacteroidetes* (Ley et al., 2005, 2006; Abdallah Ismail et al., 2011; Koliada et al., 2017). Given that marmots are obligate hibernators and fat accumulation is an indicator of good body condition and directly related to fitness in the wild (Haramis et al., 1986; Tammaru et al., 1996; Merilä & Svensson, 1997; Festa-Bianchet et al., 1997), the gut microbiome composition of marmots appears to favor weight gain that should confer greater survival and reproductive success (Armitage, 1991; Armitage, 1998). Although our results show that relative abundance of *Firmicutes* and *Bacteroidetes* is associated with mass gain rates in marmots, these results are only correlative. Further, analyses at the family level indicates selected taxa are likely involved in mass accumulation. Experimental studies similar to Stevenson, Buck, & Duddleston (2014), Stevenson, Duddleston, & Buck (2014), and Sommer et al. (2016), or multilevel structural equation modelling (Shipley, 2009) are needed to confirm causation, and to identify which bacteria are directly involved in increased mass gain rates and end of season mass gain in yellow bellied marmots. Additionally, investigating modes of microbial transmission, whether it be through social interactions, diet, or the environment could help determine why certain microbe species would be present in some individuals and not others.

The host-microbiota symbiosis is likely an important component to the hibernation phenotype. Overall, understanding the role gut microbes play in life history traits will be valuable for long-term research studies that aim to understand the implications for individual fitness. This study is novel because the majority of studies that examine the relationship between hosts and their resident microbes have been limited to humans, lab reared animals, and captive animals. Thus, studies which examine natural variation in the wild are needed if we are to understand how important these effects are in nature, and if we are to conduct robust comparative studies (Hird, 2017). Marmots are an excellent species to study to understand how microbes may impact fitness because they live in harsh, seasonal environments, however, studies on other species are needed. Gaining insight on how animals are affected by their resident microbes can help us understand the unique adaptations to harsh conditions and a changing dietary landscape and will provide essential information to future conservation and management (Mueller et al., 2020), and applications for human and animal health. Symbiotic microbes are important to animals and humans alike, and investigating this relationship in an evolutionary context is of great interest and importance.

